# Sympathetic innervation directs Ankyrin G, voltage-gated Na_v_ and KCNQ1 channels to the neuro-cardiac junctions

**DOI:** 10.1101/852129

**Authors:** Olga G. Shcherbakova

## Abstract

The sympathetic nervous system plays a central role in the cardiovascular response to acute stress by increasing heart rate and contractility. Despite the importance of functional connections between sympathetic nerves and cardiac myocytes, very little is known about the structural and functional organization of the neuro-cardiac synapses. Earlier, we demonstrated that specialized signaling domains are organized in cardiac myocytes at sites of contact with sympathetic neurons^2^. In the present study, we addressed the question if sympathetic innervation may affect localization of the cardiac ion channels. We have found that ankyrin G, scaffold protein involved in targeting ion channels and transporters, is localized at the postsynaptic sites in the cardiac myocytes innervated by sympathetic neurons. Consistent with the roles of ankyrin G in targeting Na_v_ channels to specific domains in neurons and cardiac myocytes, we have observed an increased density of Na_v_ channels at the neuro-cardiac junctions in co-cultures of cardiac myocytes and sympathetic neurons. We have also found that KCNQ1 channels are enriched at that sites. The increased density of Na_v_ and KCNQ1 channels at the sites of sympathetic innervation is likely to play a role in regulation of the cardiac excitability by sympathetic nervous system.

## Main text

Cardiac performance is regulated by neural inputs to the heart from the sympathetic and parasympathetic nervous systems^1^. According to the existing model, sympathetic neurons form *en passant* synapses with cardiac myocytes. We and other authors have demonstrated direct coupling between neurotransmitter release sites and cardiomyocytes membranes^2,3^. We have shown that the myocyte membrane develop into specialized zones that surround contacting axons and contain accumulations of the scaffold proteins SAP97 and AKAP79/150 but are deficient in caveolin-3. The β 1 ARs are enriched within these zones, whereas β 2 ARs are excluded from them after stimulation of neuronal activity^2^. Therefore, it would be of interest to further explore the molecular organization of the neuro-cardiac junctions.

Ankyrin polypeptides play critical roles in ion channels and transporters targeting to specific cellular compartments in neurons and cardiac myocytes. Ankyrin G is a key organizer of the axon initial segments in neurons since it controls the localization of membrane-associated proteins such as K_v_ and Na_v_ channels as well as the cell adhesion molecules NF186 and NrCAM ^4-6^.

In the cardiac myocytes, ankyrin G localizes at T-tubule membranes and intercalated disks. The intercalated disks serve to connect individual cardiac myocytes to work as a single functional organ or syncytium^7^. Based on the localization of ankyrin G to critical sites in cardiac myocytes, we proposed that ankyrin G might be localized at another important compartment – junctions of cardiac myocytes with sympathetic neurons.

We have tested this in the co-cultures of cardiac myocytes and sympathetic ganglion neurons (SGN) by immunostaining with antibodies to ankyrin G and tyrosine hydroxylase. Indeed, we observed that ankyrin G accumulates at the cardiac myocytes membranes at the sites of contact of cardiac myocytes and SGN (Fig.1). The fluorescence intensity for the Ankyrin G immunostaining is about twice higher at the neuro-cardiac junctions than on the non-innervated cardiac myocytes membranes (Fig.1).

**Figure 1.**
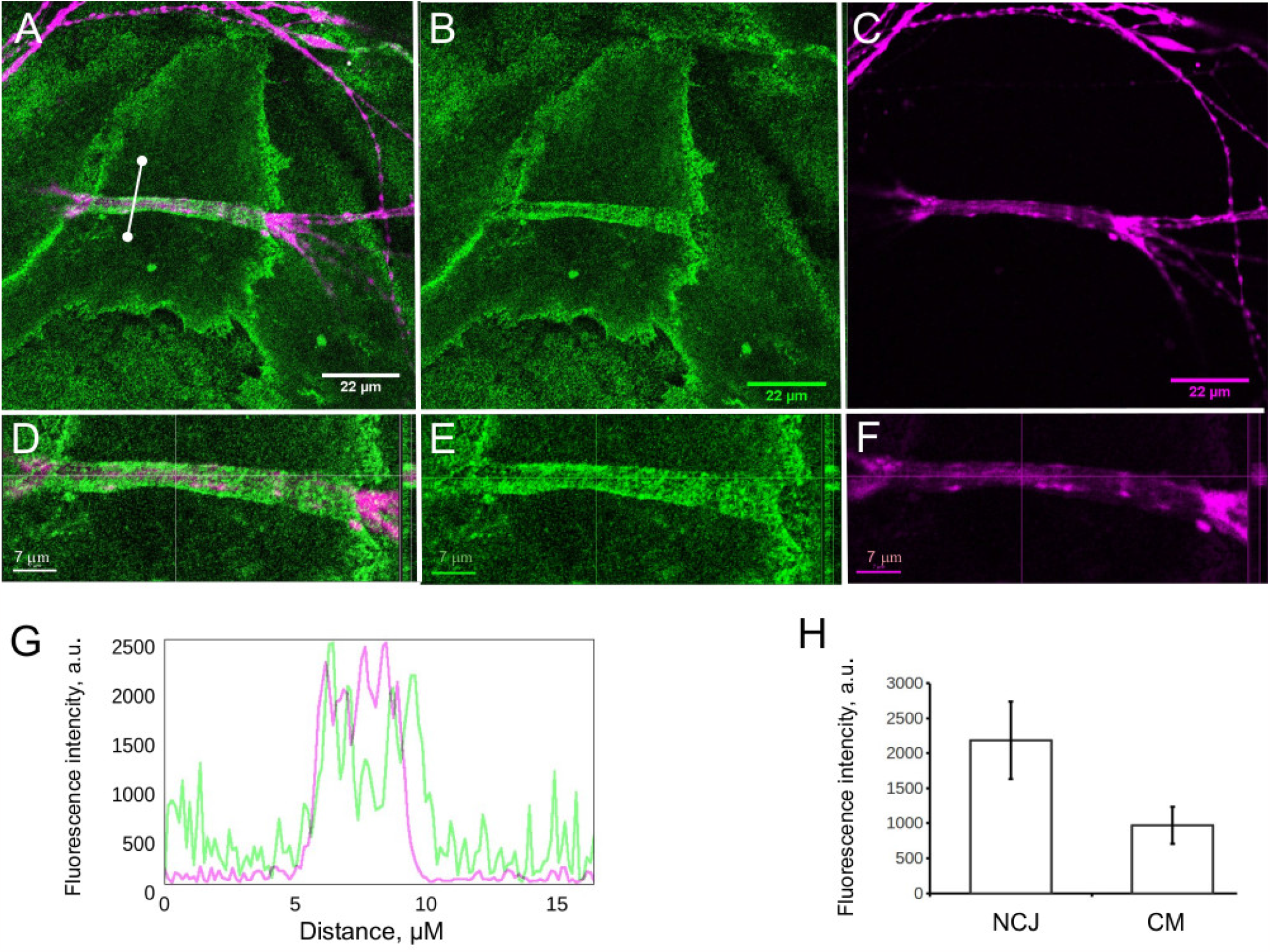
Ankyrin G is localized to the neuro-cardiac junctions. A-C. Co-culture of cardiac myocytes and sympathetic neurons (day 5 in vitro) immunostained with antibodies to Tyrosine Hydroxylase (magenta) and Ankyrin G (green). Confocal imaging, x63, single slice. D-F. 3D reconstruction of image A, an enlarged fragment corresponding to the neuro-cardiac junction (NCJ). Please note the cross-section view of the neuro-cardiac junction on the right side of the image. G. Fluorescence intensity plot along the selected straight line crossing the NCJ. H. Chart for the fluorescence intensities of the Ankyrin G immunostaining.Fluorescence intensities were quantified for the 20 ROI chosen at the neuro-cardiac junctions and 20 ROI at the cardiac myocytes surfaces as mean gray values followed by the paired T-test analysis. Measurements were performed in arbitrary units of the direct scale. Statistic comparisons were performed with a paired t-test. Error bars represent SD. ***, P < 0.001.

It is well established that Ankyrin G interacts with Na_v_ channels and retains them at the excitable domains of membranes in neurons and in cardiac myocytes ^4-8^. In cardiac myocytes ankyrin-G associates with Na_v_1.5, the primary ion channel responsible for the upstroke of the cardiac action potential. It has been demonstrated that the mutation in the Na_v_1.5 channel that abolishes the binding of Na_v_1.5 to ankyrin-G and prevents accumulation of Na_v_1.5 at cell surface sites in ventricular cardiomyocytes, cause Brugada syndrome, which is a severe form of arrhythmia leading to sudden death^9^. Consistent with the roles of ankyrin G in targeting Na_v_ channels, we have observed an increased density of Na_v_ channels using pan-Na_v_ channel antibodies (Fig.2).

**Figure 2.**
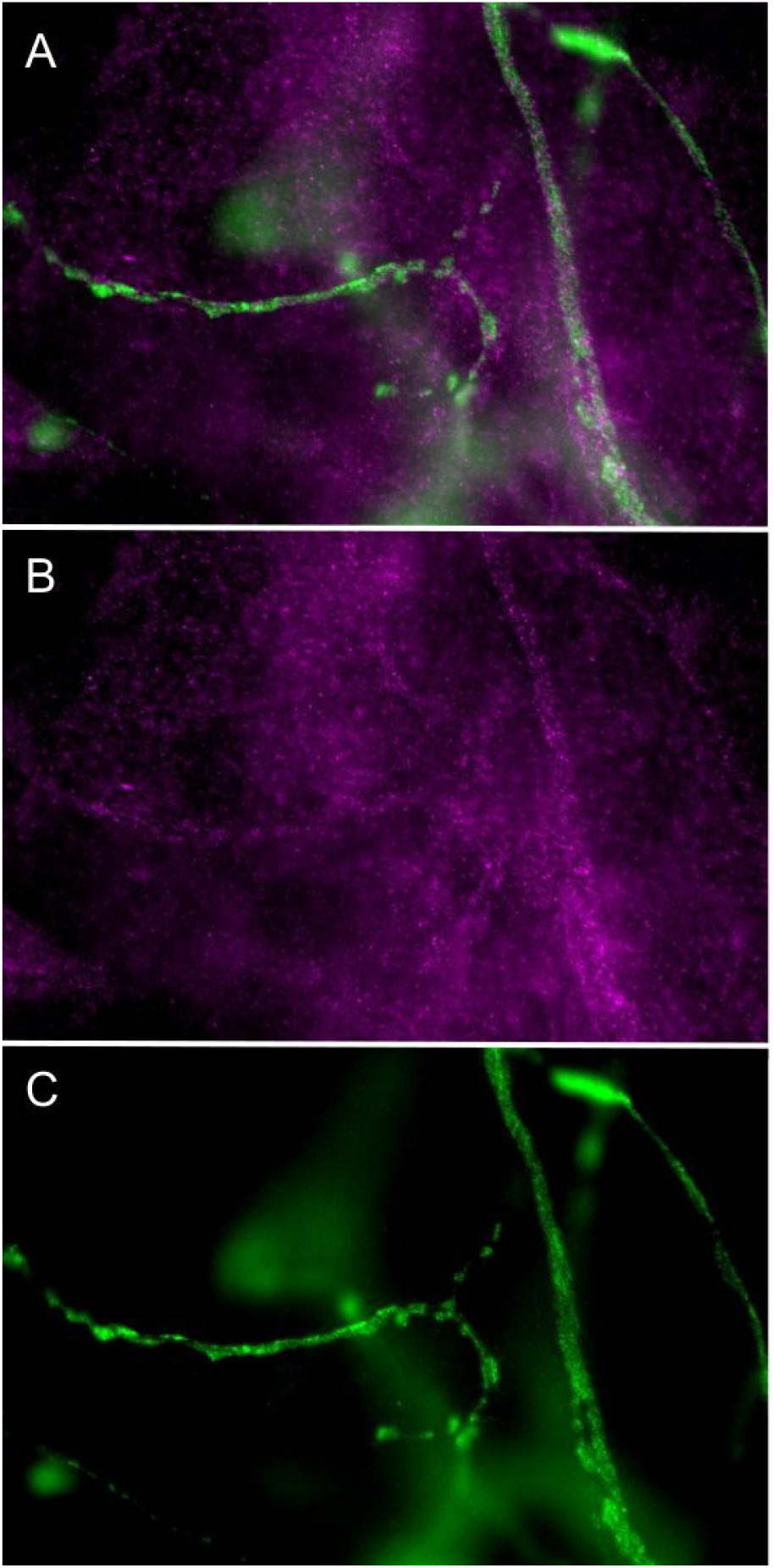
Na_v_ channels are present at the neuro-cardiac junctions. A. Co-culture of cardiac myocytes and sympathetic neurons (DIV 5) I immunostained with antibodies to Tyrosine Hydroxylase (green) and pan-Na_v_ (magenta). Epi-fluorescent image, x63. B. Immunostaining for pan-Na_v_ antibodies (red channel). C.Tyrosine Hydroxylase immunostaining (green channel).

The cardiac action potential is formed by the finely balanced activity of multiple ion channels and transporters that generate sequential events of depolarization and repolarization. In cardiac myocytes, repolarization is primarily achieved by I_ks_, mediated by KCNQ1-KCNE1 and possibly other KCNQ1-KCNE combinations^10, 11^. The input of the sympathetic nervous system, mediated by β-adrenergic receptor activation, increases the slow outward potassium ion current (I_Ks_) to accelerate repolarization, shorten the cardiac APD and increase cardiac contractility^12,13^. Thus, we have tested localization of the KCNQ1 channel at the neuro-cardiac junction. Indeed, we have observed an increased density of KCNQ1 channels at the sites of sympathetic innervation in cardiac myocytes (Fig.3).

**Figure 3.**
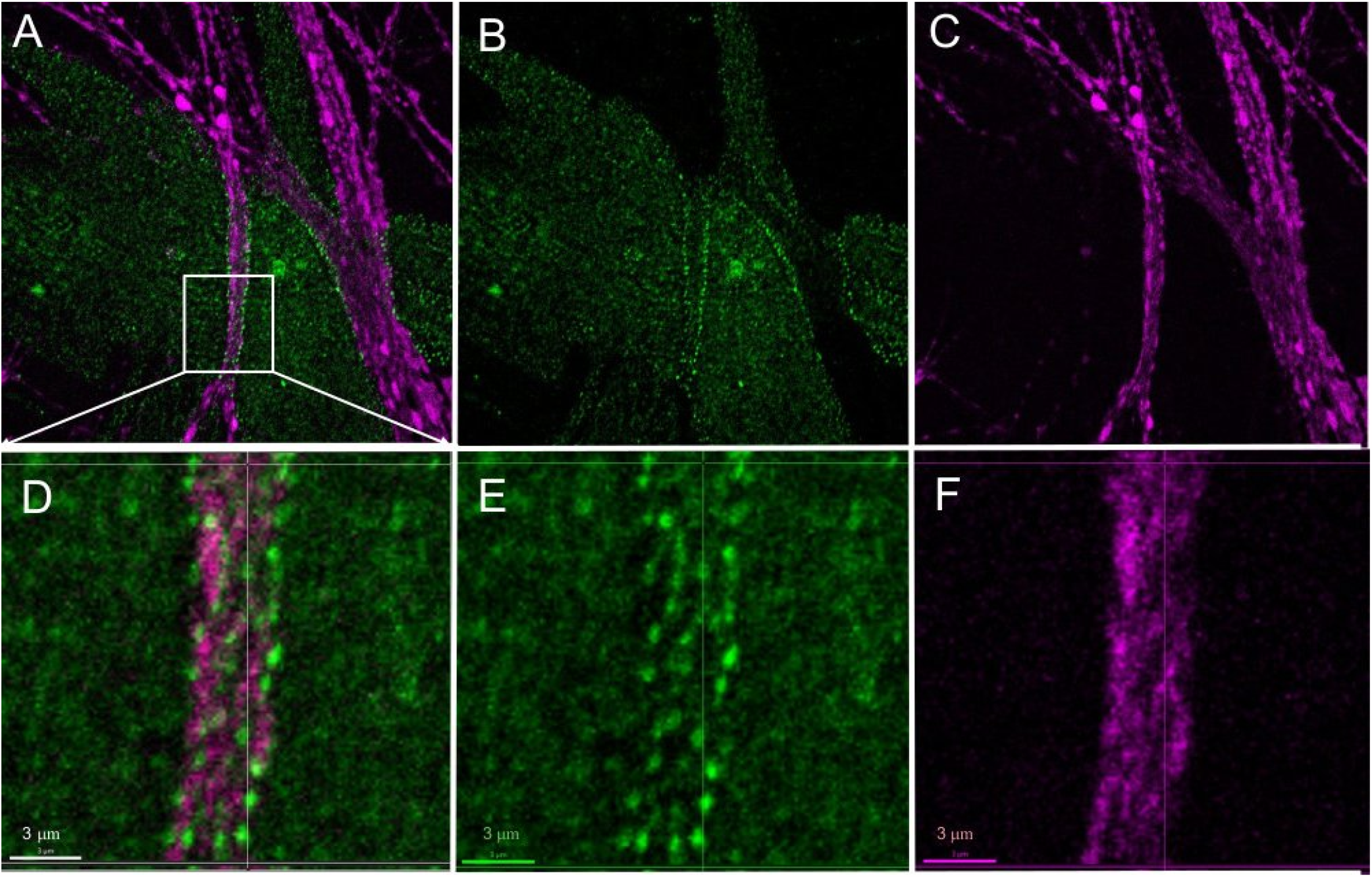
KCNQ1 channels are enriched at the neuro-cardiac junctions. A-C. Co-culture of cardiac myocytes and sympathetic neurons (DIV 5) immunostained with antibodies to Tyrosine Hydroxylase (magenta) and KCNQ1 (green). Confocalimaging, x63, single slice. D-F. 3D reconstruction of image A, an enlarged fragment corresponding to the neuro-cardiac junction.

## Discussion

Localization and targeting of the cardiac ion channels are critically important for efficient action potential initialization and propagation through the myocardium. We have found ankyrin G as well as an increased density of Na_v_ channels at the sites of sympathetic innervation in cardiac myocytes. It was demonstrated that the condition ankyrin G knockout mice display bradycardia and arrhythmia associated with catecholaminergic stress^7^. Taken together with our results, this suggests that local modulation of Na_v_ channels by sympathetic input may play an important role in the proper transmission of the catecholaminergic signals to the heart.

In human ventricular myocytes, KCNQ1-KCNE1 channels generate I_Ks_, a slowly activating K^+^ current that is important for timely myocyte repolarization. Mutations in two human I_Ks_ channel subunits, hKCNQ1 and hKCNE1, prolong action potential duration (APD), and cause inherited cardiac arrhythmias known as LQTS (long QT syndrome)^12,13^.

It is worth noting that the localization of KCNQ1 and Na_v_ channels is not exclusive to the neuro-cardiac junctions: they are enriched at these sites, but they are also expressed over the cardiac myocytes plasma membranes. This may indicate a compartmentalized regulation of these channels in the heart. For instance, it has been shown that there are two populations of K_v_4 channels responsible for I_to_ current in cardiac myocytes, one localized in caveolae and the other localized in flat rafts. Interestingly, only the population with K_v_4 channels located in caveolae can be regulated by α1-AR stimulation and its responsiveness of the K_v_4 channels to catecholaminergic stimulation depends on scaffold protein AKAP100^14^.

Variations in channels density and surface expression are known to contribute to the modulations of particular currents^15^. Therefore, increased density of ion channels at the sites of sympathetic innervation may provide a means for fine-tuning of the particular currents by neuronal input. The functional significance of localization of Na_v_ and KCNQ1 channels to the neuro-cardiac sympathetic junctions is yet to be determined.

## Acknowledgments

I am grateful to Dr. Brian K.Kobilka (Stanford University) for the opportunity to conduct the experiments in his lab and to Dr. Peter J. Mohler and Dr.Thomas Hund (The Ohio State University) for kindly providing anti-ankyrin G and anti-KCNQ1 antibodies, helpful discussion, and critical reading of the manuscript.

This work was supported by funding from the National Institutes of Health grant 1R01 HL71078-01 (to B.K. Kobilka) (experimental work) and by Russian Federation Federal budget program № AAAA-A19-119091890069-7 for Petersburg Nuclear Physics Institute named by B.P. Konstantinov of National Research Centre «Kurchatov Institute» (writing the manuscript).

## Methods

### Co-culture of sympathetic ganglion neurons (SGN) and neonatal mouse ventricular myocytes

SGN were isolated from the cervical ganglia of newborn mouse pups by treating ganglia with collagenase type 1A-S (Sigma-Aldrich) and trypsin T XI (Sigma-Aldrich) followed by trituration. All procedures met the guidelines of the National Institutes of Health Guide for the Care and Use of Laboratory Animals and were approved by the Institutional Animal Care and Use Committee at Stanford University. Neurons were plated on coverslips coated with laminin (Sigma-Aldrich) for immunocytochemistry as described previously^2^. Spontaneously beating neonatal cardiac myocytes were prepared from the hearts of newborn mouse pups as described previously^16^ and were added to already plated SGNs on the same day. After culturing for 24 h, co-cultures were treated with 1 μM cytosine arabinoside (Sigma-Aldrich) for 24 h to inhibit fibroblasts growth. Co-cultures were maintained in Leibovitz’s L-15 medium supplemented with Nu serum (BD Biosciences), NGF (Invitrogen), and ITS liquid media supplement (Sigma-Aldrich). After cytosine arabinoside treatment, media were changed every 3 d as previously described.

### Immunofluorescence microscopy

Co-cultures were fixed by adding PBS (Mediatech, Inc.) containing 8% PFA directly to the culturing media to achieve a final PFA concentration of 4%. Cells were permeabilized with 1% BSA solution in PBS containing 0.2% Triton X-100. Cells were then stained with the desired antibody. The antibodies used were as follows: anti - tyrosine hydroxylase (Transduction Laboratories, mouse monoclonal, 1:100), anti-Ankyrin G and anti-KCNQ1 (kind gift of Dr. Mohler^9^, rabbit polyclonal, 1:800); anti-tyrosine hydroxylase (Chemicon, rabbit polyclonal, 1:500), and anti-pan Na_v_ (MeuroMab, mouse monoclonal; 1:200).

The primary antibodies were detected with corresponding AlexaFluor594-conjugated goat anti–mouse IgG (1:1,000; Invitrogen) and AlexaFluor488 goat anti–rabbit IgG (1:1,000; Invitrogen). The slices for imaging were mounted with Vectashield mounting media(Vector Laboratories). The images were acquired at room temperature on an imaging microscope (Axioplan 2; Carl Zeiss MicroImaging, Inc.) using a plan-Apochromat 63X 1.40 NA oil lens (Carl Zeiss MicroImaging, Inc.), a camera (RTE/CCD-1300-Y/HS; Roper Scientific), and IPLab software (BD Biosciences).

Confocal images were acquired using a confocal laser-scanning microscope (LSM510; Carl Zeiss MicroImaging, Inc.) using Argon and He/Ne lasers and a plan-Apo 63X 1.4 NA or plan-Apo 100x 1.1 NA oil lenses, and images were analyzed by ImageJ and Imaris software. Immunostaining experiments using sets of six separate coverslips were repeated at least three times.

